# A PEROXO-Tag enables rapid isolation of peroxisomes from human cells

**DOI:** 10.1101/2020.03.10.984948

**Authors:** G. Jordan Ray, Elizabeth A. Boydston, Emily Shortt, Gregory A. Wyant, Sebastian Lourido, Walter W. Chen, David M. Sabatini

## Abstract

Peroxisomes are metabolic organelles that perform a diverse array of critical functions in human physiology. Traditional isolation methods for peroxisomes can take more than one hour to complete and can be laborious to implement. To address this, we have now extended our prior work on rapid organellar isolation to peroxisomes via the development of a peroxisomally-localized 3XHA epitope tag (“PEROXO-Tag”) and associated immunoprecipitation (“PEROXO-IP”) workflow. Our PEROXO-IP workflow has excellent reproducibility, is easy to implement, and achieves highly rapid (~10 minutes post-homogenization) and specific isolation of human peroxisomes, which we characterize here via proteomic profiling. By offering speed, specificity, reproducibility, and ease of use, the PEROXO-IP workflow should facilitate studies on the biology of peroxisomes.

## INTRODUCTION

Peroxisomes are membrane-bound organelles that perform a diverse array of biological functions in human cells. Defects in peroxisomal biogenesis or associated enzymatic functions can lead to devastating pathologies that manifest early in life (1). With regards to the study of peroxisomes, direct interrogation of isolated peroxisomes can provide insights into peroxisomal physiology beyond that achieved using whole-cell analyses.

We have previously developed multiple workflows that utilize 3XHA epitope tags localized to either mitochondria (2–4) or lysosomes (5) to enable rapid immunocapture of the respective subcellular compartment. These methods have been successfully used in conjunction with downstream metabolomic analyses to study the dynamics of mitochondria (2, 4) and with metabolomic and proteomic analyses to study the behavior of lysosomes (5–7) and identify changes that were not readily visible with whole-cell or whole-tissue analyses. Historically, isolating peroxisomes has been one of the more challenging enterprises in organellar purification because peroxisomes can be fragile and share similar biophysical properties (e.g., density) with other organelles, such as mitochondria and lysosomes (8). In addition, the vast majority of isolation methods for peroxisomes have relied on workflows that can be lengthy and laborious to implement. Indeed, many of the common methods for isolating peroxisomes involve density-gradient centrifugation workflows that can last more than one hour from the time of homogenization (8–11), increasing the likelihood for distortion of the native organellar profile (e.g., losing associated peripheral membrane proteins) and damage to the organelle itself. Alternative approaches, such as magnetic-bead-based affinity purification of peroxisomes via endogenous ABCD3, a peroxisomal membrane protein (12, 13), or workflows utilizing zonal free flow electrophoresis (14) or immune free flow electrophoresis (15) all can take more than one hour from the time of homogenization as well.

To address these issues, we have extended our original epitope-tagging approach (2, 5) to peroxisomes and generated a “PEROXO-Tag” that allows for rapid (~10 min post-homogenization), specific, and facile isolation of peroxisomes in a highly reproducible manner. The PEROXO-Tag is a chimeric protein consisting of three HA epitopes attached to the N-terminus of monomeric EGFP and a segment of PEX26 fused to the EGFP C-terminus, which allows for localization to and insertion into the peroxisomal membrane. Leveraging the high-affinity, high-specificity interaction between the tandem HA epitopes of the PEROXO-Tag and the cognate anti-HA antibody, we show that this PEROXO-Tag and the associated immunoprecipitation (“PEROXO-IP”) workflow can be used on human cells for the rapid immunoisolation of peroxisomes, which we characterize here via proteomic analysis. Our PEROXO-IP workflow also exhibits high reproducibility and allows for straightforward isolation of peroxisomes from cells expressing the PEROXO-Tag. Taken together, this work extends our previous efforts for developing a tool kit of epitope-tag-based handles for the rapid and specific isolation of organelles to peroxisomes.

## RESULTS

To enable rapid isolation of peroxisomes from human cells, we generated the PEROXO-Tag, a chimeric protein in which three HA epitopes are attached to the N-terminus of monomeric EGFP and amino acids 237-305 of human PEX26 are fused to the EGFP C-terminus, allowing for localization to and insertion into the peroxisomal membrane (3XHA-EGFP-PEX26 or HA-PEROXO). As a control, we also designed a counterpart protein in which the three HA epitopes were exchanged for three Myc epitopes (3XMyc-EGFP-PEX26 or Control-PEROXO) (Fig. 1*A*). Amino acids 237-305 of human PEX26 have been used previously to localize a protein to peroxisomes (16). We chose to use this segment of PEX26, rather than full-length PEX26, to minimize unintended biological effects from exogenous expression of the *Control-PEROXO* and *HA-PEROXO* genes. Indeed, human PEX26 is a peroxin that interacts with the PEX1-PEX6 complex, as well as PEX14, and it has been reported previously that amino acids 13-48 and 78-85 in the N-terminal region of PEX26 are required for the interaction with the PEX1-PEX6 complex and PEX14, respectively (17).

**Fig. 1.**
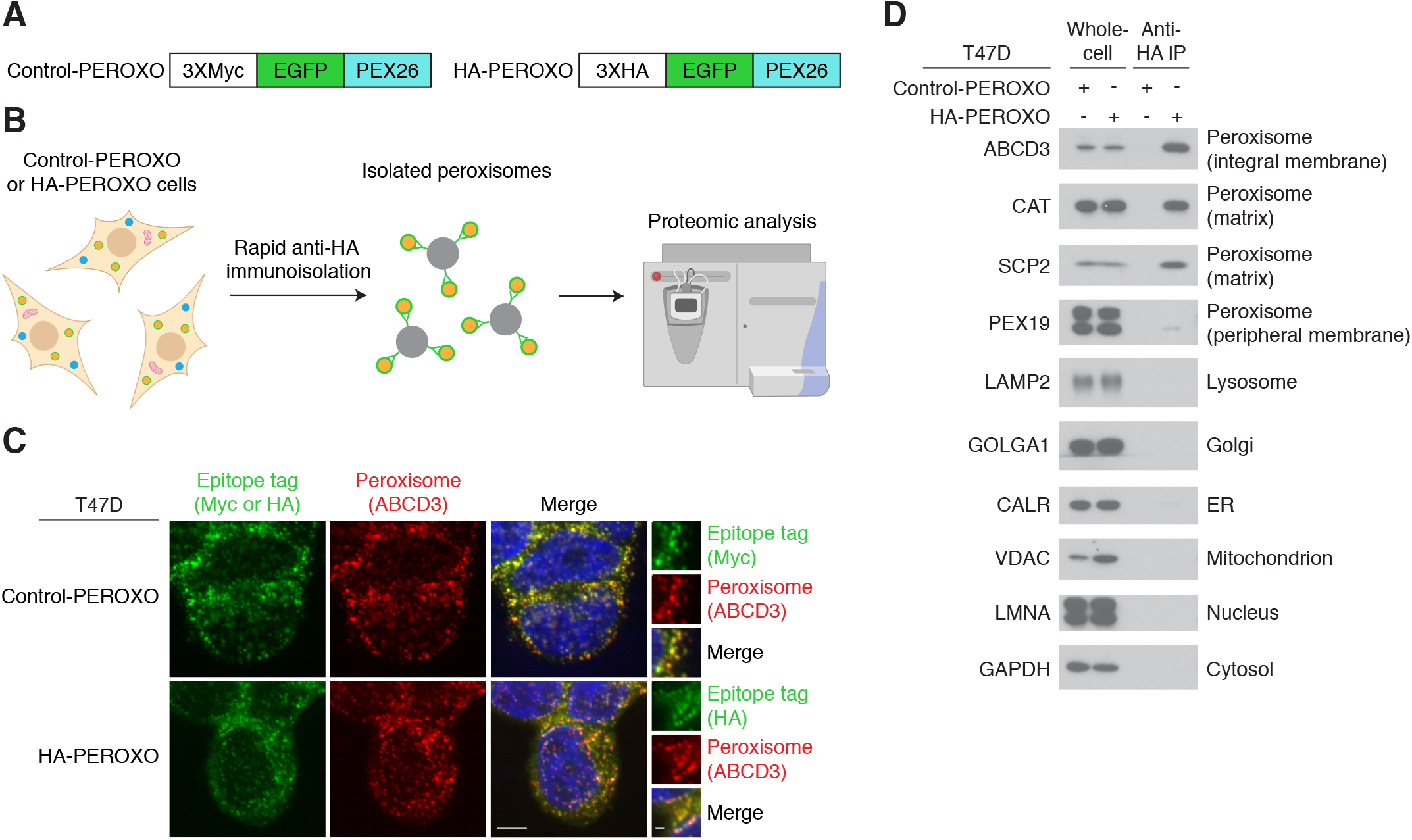
A workflow for the rapid and specific isolation of peroxisomes from human cells. (*A*) The design of the 3XMyc-EGFP-PEX26 (Control-PEROXO) and 3XHA-EGFP-PEX26 (HA-PEROXO) proteins. EGFP, monomeric EGFP; PEX26, amino acids 237-305 of human PEX26. (*B*) Schematic of the PEROXO-IP workflow, which allows for rapid (~10 min once cells are homogenized) and specific isolation of peroxisomes from human cells for downstream profiling via approaches such as proteomic analysis. The immunocapture workflow used here is generally similar to that reported previously (3). (*C*) Immunofluorescence of the indicated T47D cell line. Representative images with epitope-tag (identified by Myc-positive or HA-positive structures) (green), peroxisomal (identified by ABCD3-positive structures) (red), and nuclear (blue) signals are shown. Staining of epitope tags was done despite the presence of EGFP in the respective proteins because cells are sorted for low EGFP levels in our methodology, thus rendering the EGFP signal weak. Insets represent zoomed-in fields. The scales bars in the large and small images represent 5 μm and 1 μm, respectively. (*D*) Immunoblot analysis of T47D whole cells and immunoprecipitates from the PEROXO-IP workflow. The names of the protein markers used and the corresponding subcellular compartments appear to the left and right of the immunoblots, respectively. Golgi, Golgi complex; ER, endoplasmic reticulum.

In our PEROXO-IP workflow, we utilize cells lentivirally transduced with a 3XMyc-EGFP-PEX26 construct (Control-PEROXO cells) or a 3XHA-EGFP-PEX26 construct (HA-PEROXO cells). Both Control-PEROXO and HA-PEROXO cells are sorted for low EGFP levels via FACS to reduce the chances of unwanted effects that might be secondary to excessive amounts of the Control-PEROXO or HA-PEROXO proteins, such as mislocalization of the epitope-tagged proteins and perturbations of native cell biology. Utilizing the PEROXO-IP workflow, we are able to achieve rapid peroxisomal isolation in ~10 min post-homogenization using a methodology that is generally similar to that described previously (3). This is significantly faster than the vast majority of prior isolation methods, which can require more than one hour to isolate peroxisomes after the starting material has been homogenized. In addition, the peroxisomal isolation buffer used in this workflow (i.e., “KPBS”), which we had developed initially for rapid isolation of mitochondria (2), is compatible with mass spectrometry, thus allowing for downstream mass-spectrometric analyses, such as proteomics (Fig. 1*B*).

Importantly, we found that our PEROXO-IP workflow behaved as anticipated when evaluated via orthogonal methods. As part of our initial characterization of the PEROXO-IP methodology, we utilized a standard cultured human cell line, T47D. Immunofluorescence-based examination revealed co-localization of the Control-PEROXO and HA-PEROXO proteins with the peroxisomal marker ABCD3 (Fig. 1*C*). Importantly, immunoblot analysis of the immunoprecipitates using Control-PEROXO cells (Control-PEROXO IP) and HA-PEROXO cells (HA-PEROXO IP) revealed enrichment of peroxisomes relative to a variety of other organelles and minimal organellar contamination of the Control-PEROXO IP (Fig. 1*D*), demonstrating that our PEROXO-IP workflow can achieve rapid and specific isolation of peroxisomes. We assessed peroxisomes using a variety of different markers, namely ABCD3, CAT, SCP2, and PEX19. The form of SCP2 we examined is the ~14 kDa product that results from transcription of a downstream promoter for the *SCP2* gene and has been used previously to better assess the integrity of isolated peroxisomes (18). PEX19 is a peripheral membrane protein that shuttles between the cytosol and surface of peroxisomes; the percentage of the total cellular population of PEX19 that is actually on peroxisomes can be variable (19), but we did observe the presence of the protein in our peroxisomal isolates from both T47D cells (Fig. 1*D*) and HEK-293T cells (Fig. 3*A*), suggesting that the workflow can even capture non-integral membrane proteins that interact with the peroxisome on a more transient basis, which may be a reflection of the rapidity of the method. It is interesting that in T47D cells, CAT does not exhibit similar patterns of enrichment as ABCD3 and SCP2, which is unlikely to be a reflection of CAT loss from damaged, isolated peroxisomes as SCP2, a smaller matrix protein, does not behave similarly. Instead, the pattern of enrichment of CAT may reflect the presence of some CAT in organelles outside of peroxisomes (20).

We next examined whether introduction of the Control-PEROXO or HA-PEROXO proteins perturbed native features of peroxisomes, such as levels of known peroxisomal proteins or the morphology and distribution of peroxisomes. Using immunoblot analysis of whole cells, we did not observe any noticeable difference in the levels of standard peroxisomal markers such as ABCD3 and CAT, or the peroxisomal proteins PEX19, SCP2, and endogenous PEX26 in Control-PEROXO or HA-PEROXO cells versus those transduced with the corresponding empty vector (Fig. 2*A*). In addition, immunofluorescence-based examination of peroxisomes revealed no clear changes to peroxisomal morphology or distribution (Fig. 2*B*).

**Fig. 2.**
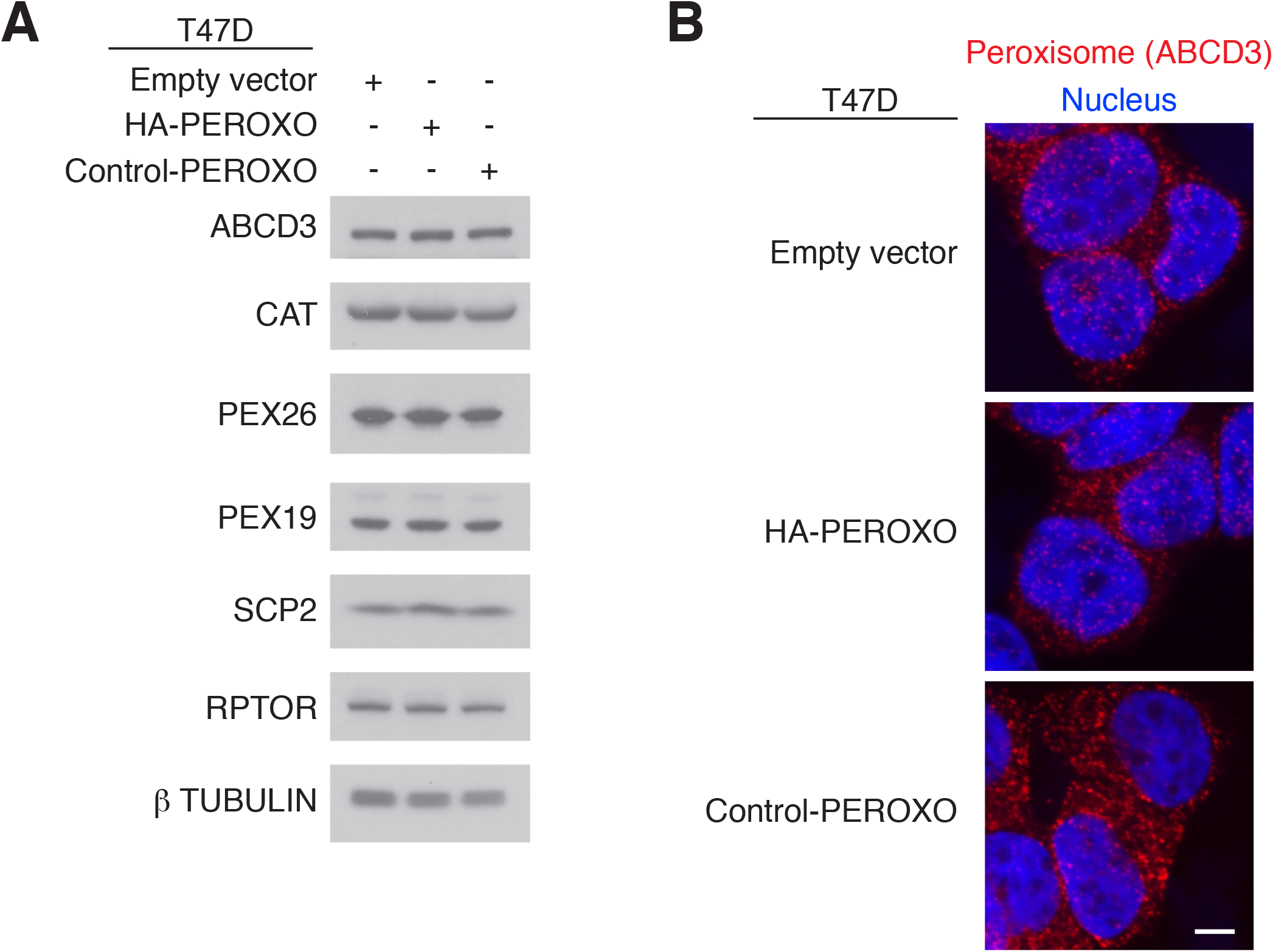
Examination of the effects of the Control-PEROXO and HA-PEROXO proteins on peroxisomes. (*A*) Whole-cell immunoblot analysis of the indicated T47D cell line. The names of the proteins appear to the left. RPTOR and β TUBULIN are loading controls. (*B*) Immunofluorescence of the indicated T47D cell line. Representative images with peroxisomal (identified by ABCD3-positive structures) (red) and nuclear (blue) signals are shown. The scale bar represents 5 μm.

To better characterize the peroxisomes isolated using our PEROXO-IP workflow, we analyzed whole-cell and immunoprecipitate (IP) material using quantitative proteomics and data-independent acquisition. To increase yield for the proteomic work, we scaled up the input and transitioned the PEROXO-IP workflow to HEK-293T cells. We confirmed that our PEROXO-IP workflow still resulted in substantial enrichment of peroxisomes relative to other subcellular compartments under these conditions (Fig. 3*A*). The PEROXO-IP workflow also demonstrated excellent reproducibility across either Control-PEROXO IP or HA-PEROXO IP biological replicates, as evidenced by the overlap of measured abundances in scatter plots and by correlational analysis exhibiting Spearman rs values > 0.84 in all comparisons (Fig. 3*B*). Of the 1441 proteins in our primary proteomic data, 288 proteins display greater signal in the HA-PEROXO IP samples than in the Control-PEROXO IP samples (i.e., the background) and are statistically significant after correcting for multiple hypothesis testing - these 288 proteins are considered the “top proteins” (Dataset S1). 32 of these top proteins are annotated as peroxisomal per the Gene Ontology (GO) Resource; for context, the GO Resource has a total of 141 proteins assigned to peroxisomes in humans. To account for factors such as absent or low expression of genes encoding for certain peroxisomal proteins in HEK-293T cells and/or poor behavior of certain peroxisomal proteins with regards to proteomic analysis, we examined all 1441 proteins of the primary proteomic data, which represent proteins found in IP and/or whole-cell samples, and found 45 peroxisomal proteins. We thus determined the general coverage of the PEROXO-IP workflow to be ~71% (32/45) (Dataset S1 and Fig. 3*C*). Importantly, the peroxisomal proteins ABCD3, CAT, and SCP2 are present among the top proteins, corroborating the results of our immunoblot analysis (Fig. 3 *A* and *C*). Although we could detect PEX19 in our peroxisomal isolates via immunoblotting (Fig. 3*A*), it is not present in our proteomic data (Dataset S1), which may reflect a combination of low protein abundance and/or the performance of this protein in our proteomics. Mitochondria and endoplasmic reticulum (ER) proteins can be commonly seen in density-gradient centrifugation preparations of peroxisomes (21, 22); as such, we interrogated the top proteins for signs of those two organelles, and found 182 mitochondrial proteins and 66 ER proteins per the GO assignments (Dataset S1*C*). This mitochondrial signal is potentially a result of native mitochondrial-peroxisomal contacts (23), transfer of mitochondrial material to peroxisomes via mitochondrial-derived vesicles (24, 25), and/or the contribution of mitochondria to the *de novo* synthesized pool of peroxisomes (26). Similarly, this ER signal could be present because of peroxisomal-ER contacts (23) and/or the contribution of ER to peroxisomes during *de novo* biogenesis (1, 26). Importantly though, by using an organellar enrichment metric (i.e., HA-PEROXO IP signal / HA-PEROXO whole-cell signal) for the top proteins that also behave as ideal markers (i.e., strictly annotated to only one organelle in the set of peroxisome, mitochondria, or ER), we found that our PEROXO-IP workflow enriched for peroxisomes to a significantly greater degree than for mitochondria or ER (Fig. 3*D* and Dataset S1*C*). We chose proteins only assigned to one of the three organelles to minimize the confounding effects of including proteins that could have multiple localizations. Reassuringly, the median organellar enrichment for peroxisomes is ~12.7-fold greater than for mitochondria and ~14-fold greater than for ER, which strongly corroborates the results from our immunoblot analysis (Fig. 3*A* and Dataset S1*C*) and demonstrates that, despite the presence of mitochondrial and ER proteins in the IP material, mitochondria and ER were not enriched nearly to the same degree as peroxisomes were. With regards to the 32 peroxisomal proteins belonging to the group of top proteins, we examined each of their associated functions and found a diverse range of molecular players involved in classic processes associated with peroxisomes, such as peroxins (e.g., PEX3, PEX11B, PEX13, PEX14), metabolic proteins involved in the oxidation of very long-chain fatty acids (e.g., ACOX1, HSD17B4) or synthesis of ether lipids (e.g., AGPS, GNPAT), and antioxidant defense proteins (e.g., CAT, GSTK1) (Fig. 3*E*). Taken together, these data demonstrate that our PEROXO-IP workflow can be used to rapidly isolate peroxisomes for downstream interrogation of the peroxisomal proteome.

**Fig. 3.**
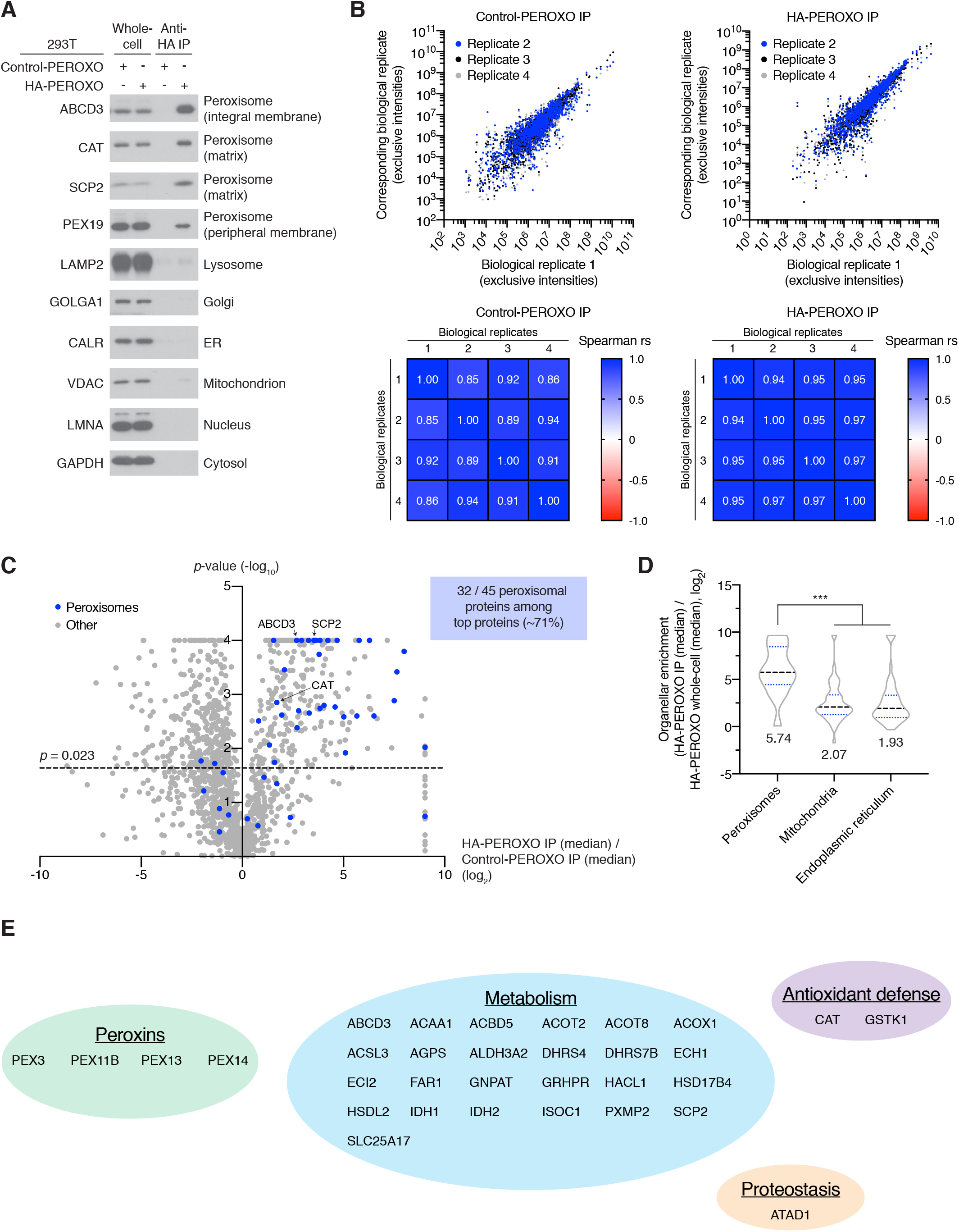
Proteomic characterization of isolated peroxisomes. (*A*) Immunoblot analysis of HEK-293T whole cells and immunoprecipitates from the PEROXO-IP workflow. The names of the protein markers used and the corresponding subcellular compartments appear to the left and right of the immunoblots, respectively. Golgi, Golgi complex; ER, endoplasmic reticulum. (*B*) Examination of reproducibility among Control-PEROXO IP and HA-PEROXO IP biological replicates subjected to proteomic interrogation. Scatter plots of the reported exclusive intensities for each protein for each biological replicate of the indicated IP group and correlation matrices demonstrating the Spearman rs value for each indicated comparison are shown. The primary proteomic data shown in Dataset S1*B* were used to generate this panel. For all correlation analyses of different replicates in either the Control-PEROXO IP group or HA-PEROXO IP group, *p* < 0.001 and all correlations were significant after application of the Benjamini-Hochberg procedure (FDR = 5%). For the purposes of plotting and analysis, only proteins with reported exclusive intensities in all four biological replicates of a given IP group were included for that IP group’s scatter plot and correlation analyses; accordingly, n = 1360 proteins for the Control-PEROXO IP group and n = 1426 proteins for the HA-PEROXO IP group. (*C*) Scatter plot of proteins identified by proteomics in the immunoprecipitates from the PEROXO-IP workflow. The primary proteomic data shown in Dataset S1*B* were used to generate this panel. The dotted line indicates where the *p*-value = 0.023; all points above this line are those that are significant after application of the Benjamini-Hochberg procedure (FDR = 5%). For the purposes of graphing, only proteins that have a value for log2(HA-PEROXO IP (median) / Control-PEROXO IP (median)) are plotted here (n = 1434 proteins). Regardless, all 45 proteins annotated as peroxisomal per the Gene Ontology (GO) Resource in our primary proteomic data (n = 1441 proteins) are plotted here and indicated in blue and, of these 45 peroxisomal proteins, 32 are among the top proteins from our analysis (i.e., have a log_2_ score > 0 and are statistically significant after application of the Benjamini-Hochberg procedure). Proteins with no peroxisomal annotation are in grey. (*D*) Violin plots of the organellar enrichments for peroxisomes, mitochondria, or endoplasmic reticulum. Top proteins shown in Dataset S1*C* were used to generate this plot, but only proteins that were localized by the GO Resource solely to the indicated organelle and not to the other two organelles were used to represent the corresponding organelle. Proteins were chosen in this way to minimize the confounding effects of including proteins that could have localization to more than one of the three organelles. Peroxisomes (n = 15 proteins), mitochondria (n = 159 proteins), endoplasmic reticulum (n = 56 proteins). Black dotted line indicates the median, blue dotted lines the quartiles, and the violin plots extend from the minimum to maximum of each data set. The actual numerical value of the median is shown below each corresponding violin plot. \*\*\**p* < 0.001. (*E*) Identities and general classifications of the 32 previously annotated peroxisomal proteins found among the top proteins. The corresponding NCBI gene symbols are shown. For all proteomic analysis in Fig. 3, see Dataset S1 for additional data and details.

## DISCUSSION

Building upon our prior work utilizing epitope tags for rapid isolation of mitochondria (2, 4) and lysosomes (5), we have now developed a PEROXO-Tag and PEROXO-IP workflow that allows for rapid (~10 min post-homogenization), specific, and facile isolation of peroxisomes from cultured human cells in a highly consistent manner. Introduction of the PEROXO-Tag into cells did not noticeably alter native features of peroxisomes, such as the whole-cell levels of various peroxisomal proteins and the morphology and distribution of peroxisomes. Proteomic analysis of peroxisomes isolated using our PEROXO-IP workflow identified proteins that are known to be peroxisomal components that participate in a diverse range of biological processes.

During the course of this project, a separate study by Xiong *et al.* was published describing the development of a rapid immunopurification scheme for mitochondria, lysosomes, and peroxisomes using a twin-strep-tag localized to the respective organelles (27). The peroxisome-localization sequence used in their work is based on PEX3, whereas ours is based on PEX26. However, compared to the study by Xiong *et al.*, our work goes into substantially greater depth characterizing our Control-PEROXO and HA-PEROXO proteins and our peroxisomal isolates. From a broader perspective, no direct comparison was done between our 3XHA-based approaches for mitochondria (2, 3) and lysosomes (5) and the twin-strep-tag-based approaches, so it is difficult to assess how the two epitope-tagging strategies generally compare. Also of note, the work by Xiong *et al.* does not report usage of a control IP, which is essential for assessing background binding of materials to beads and is concerning with respect to their metabolomic interrogation of lysosomes. Indeed, from multiple studies (2, 4–7), including the work shown here, we have found the control IP to be critical for bead-based affinity purification of organelles when combined with polar metabolomic, lipidomic, or proteomic analyses.

In summary, we believe that the PEROXO-IP workflow described here will be useful for the isolation and study of peroxisomes given the rapidity, specificity, ease of use, and reproducibility of the methodology. The ability to rapidly isolate peroxisomes can be helpful for the study of proteins by better preserving the dynamic interactions of peripheral membrane proteins with the peroxisomal membrane. Rapidity of isolation can also help preserve the native proteomic profile of peroxisomes, given that these organelles can be fragile once released from cells (8). The ease of use and speed of isolation should also facilitate scaling up studies on peroxisomes in cultured cells and can thus be useful for examining peroxisomes under multiple genetic, pharmacologic, or environmental perturbations, particularly since the workflow does not require the generation of density-gradients. In addition, translation of this methodology to an *in vivo* mammalian system should be possible using a strategy we have implemented previously for mitochondria (4) and thereby enable rapid isolation of peroxisomes from specific cell types without the need for cell sorting. We thus believe that the PEROXO-Tag and PEROXO-IP workflow described in this study will have utility for studying peroxisomes and can help elucidate the more dynamic processes associated with these organelles during various states of cellular function.

## MATERIALS AND METHODS

### Reagents and equipment

Reagents and equipment were obtained from the following sources: Gibson Assembly Master Mix (E2611), XhoI (R0146L), NotI (R0189L), AgeI HF (R3552L), and EcoRI HF (R3101L) were purchased from New England BioLabs; anti-HA magnetic beads (88836) and DynaMag Spin Magnet (12320D) from Thermo Fisher Scientific; plunger (89026-398) and glass vessel (89026-386) for homogenizing cells from VWR; Odyssey Buffer from Licor (927-40000); Vectashield containing DAPI from Vector Laboratories (H-1200); LoBind tubes from Eppendorf (022431081); and PBS from Thermo Fisher Scientific (10-010-023). KPBS was prepared as described previously (3) and was comprised of 136 mM KCl, 10 mM KH_2_PO_4_, pH 7.25 in Optima, LC/MS-grade water and was prepared using clean equipment and glassware. The pH of KPBS was adjusted with KOH.

### Antibodies

Antibodies were obtained from the following sources and used with the indicated dilutions: rabbit anti-ABCD3 (1:1000 for immunoblotting, 1:400 for immunofluorescence) was purchased from Abcam (ab3421); rabbit anti-CAT (1:1000) from CST (12980S); rabbit anti-SCP2 (1:500) from Proteintech (23006-1-AP); mouse anti-PEX26 (1:500) from Santa Cruz Biotechnology (sc-376817); rabbit anti-PEX19 (1:500) from Abcam (ab137072); mouse anti-LAMP2 (1:1000) from Santa Cruz Biotechnology (sc-18822); rabbit anti-GOLGA1 (1:1000) from CST (13192); rabbit anti-CALR (1:1000) from CST (12238); rabbit anti-VDAC (1:1000) from CST (4661); mouse anti-LMNA (1:1000) from CST (4777); rabbit anti-GAPDH (1:1000) from CST (2118); rabbit anti-RPTOR (1:1000) from EMD (09-217); mouse anti-β TUBULIN (1:1000) from Santa Cruz Biotechnology (sc-365791); mouse anti-HA (1:400) from CST (2367S); mouse anti-Myc (1:400) from CST (2276S); 594 donkey anti-rabbit (1:400) from Invitrogen (A32754); 488 donkey anti-mouse (1:400) from Invitrogen (A32766); anti-rabbit IgG (1:3000) from CST (7074); and anti-mouse IgG (1:3000) from CST (7076).

### Cloning and generation of lentiviral constructs

pLJC5 with 3XMyc-EGFP-PEX26 (Control-PEROXO) and pLJC5 with 3XHA-EGFP-PEX26 (HA-PEROXO) lentiviral constructs were designed as follows. The 3XMyc-EGFP and 3XHA-EGFP portions from our prior 3XMyc-EGFP-OMP25 and 3XHA-EGFP-OMP25 constructs (2) were used, respectively, with the one modification being introduction of an A206K mutation into the EGFP to monomerize it (28). The portion of human PEX26 that was used in the Control-PEROXO and HA-PEROXO proteins corresponds to amino acids 237-305 of human PEX26 and has been used previously to localize a protein to peroxisomes (16). We chose to use this segment of PEX26, rather than full-length PEX26, to minimize unintended biological effects from exogenous expression of the *Control-PEROXO* and *HA-PEROXO* genes. Indeed, human PEX26 is a peroxin that interacts with the PEX1-PEX6 complex, as well as PEX14, and it has been reported previously that amino acids 13-48 and 78-85 in the N-terminal region of PEX26 are required for the interaction with the PEX1-PEX6 complex and PEX14, respectively (17).

In terms of the details of how the constructs were generated, the 3XHA-EGFP-PEX26 sequence was initially cloned into a pMXs-IRES-blasticidin retroviral vector (CellBiolabs, RTV-016) to generate the pMXs-3XHA-EGFP-PEX26 plasmid via digestion with XhoI and NotI, Gibson Assembly, Gibson Assembly Master Mix, and the gBlocks “pMXs_3XHA-EGFP-PEX26_block1” and “pMXs_3XHA-EGFP-PEX26_block2” from Integrated DNA Technologies (see below).

pMXs_3XHA-EGFP-PEX26_block1:

TACGGGAATTCCTGCAGGCCTCGAGGCCACCATGTATCCCTATGACGTGCCTGATT ACGCCGGCACAGGATCCTACCCCTATGATGTGCCTGACTACGCTGGCAGCGCCGG ATACCCTTATGATGTGCCTGATTATGCTGGAGGGAGCGGCGTGAGCAAGGGCGAG GAGCTGTTCACCGGGGTGGTGCCCATCCTGGTCGAGCTGGACGGCGACGTAAAC GGCCACAAGTTCAGCGTGTCCGGCGAGGGCGAGGGCGATGCCACCTACGGCAAG CTGACCCTGAAGTTCATCTGCACCACCGGCAAGCTGCCCGTGCCCTGGCCCACCC TCGTGACCACCCTGACCTACGGCGTGCAGTGCTTCAGCCGCTACCCCGACCACAT GAAGCAGCACGACTTCTTCAAGTCCGCCATGCCCGAAGGCTACGTCCAGGAGCGC ACCATCTTCTTCAAGGACGACGGCAACTACAAGACCCGCGCCGAGGTGAAGTTCG AGGGCGACACCCTGGTGAACCGCATCGAGCTGAAGGGCATCGACTTCAAGGAGGA CGGCAACATCCTGGGGCACAAGCTGGAGTACAACTACAACAGCCACAACGTCTATA TCATGGCCGACAAGCAGAAGAACGGCATCAAGGTGAACT

pMXs_3XHA-EGFP-PEX26_block2:

ATGGCCGACAAGCAGAAGAACGGCATCAAGGTGAACTTCAAGATCCGCCACAACAT CGAGGACGGCAGCGTGCAGCTCGCCGACCACTACCAGCAGAACACCCCCATCGG CGACGGCCCCGTGCTGCTGCCCGACAACCACTACCTGAGCACCCAGTCCAAGCTG AGCAAAGACCCCAACGAGAAGCGCGATCACATGGTCCTGCTGGAGTTCGTGACCG CCGCCGGGATCACTCTCGGCATGGACGAGCTGTACAAGGGAGGGAGCGGACGCC AGCTTTGGGACTCTGCGGTGAGCCACTTCTTTTCTCTGCCCTTCAAAAAGAGTCTC CTGGCTGCCTTGATCCTCTGTCTCCTGGTGGTGAGATTTGATCCAGCTTCCCCTTC CTCCCTGCACTTCCTCTACAAGCTGGCCCAGCTCTTCCGCTGGATCCGGAAGGCT GCATTTTCTCGCCTCTACCAGCTCCGCATCCGTGACTGAGCGGCCGCTACGTAAAT TCCGCCC

We subsequently transitioned to pLJC5, a lentiviral vector that has previously been shown to afford stable, moderate expression of genes via the UBC promoter and been successfully used for rapid organellar isolation (5). The relevant oligonucleotide sequences used for transitioning to pLJC5 are shown below and were from Integrated DNA Technologies.

oGJR224 (forward primer for cloning the *3XHA-EGFP-PEX26* gene into pLJC5):

TAGCGCTACCGGTTTAATTAAGCCACCATGTATCCCTATGACGTGC

oGJR225 (reverse primer for cloning either *3XMyc-EGFP-PEX26* or *3XHA-EGFP-PEX26* genes into pLJC5):

CGAGGTCGAGAATTCTTATCAGTCACGGATGCGGAGC

oGJR226 (forward primer for cloning the *3XMyc-EGFP-PEX26* gene into pLJC5):

TAGCGCTACCGGTTTAATTAAGCCACCATGGAGCAGAAGCTGATTTCTGAGGAAGATCTGGGCACAGGATCCGAACAGAAACTGATTTCTGAGGAAGATCTGGGCAGCGCCGGAGAGCAGAAGCTGATTTCTGAAGAGGATCTGGGAGGGAGCGGCGTGAGCAAGGGCGAGGAGCT

oGJR237 (oligonucleotide used to insert into pLJC5 top strand for generating the pLJC5 empty vector):

CCGGTAAGTTCATCTGCACCACCGGCG

oGJR238 (oligonucleotide used to insert into pLJC5 bottom strand for generating the pLJC5 empty vector):

AATTCGCCGGTGGTGCAGATGAACTTA

To create pLJC5-3XHA-EGFP-PEX26, the 3XHA-EGFP-PEX26 sequence was amplified by PCR using oGJR224 and oGJR225 from the pMXs-3XHA-EGFP-PEX26 plasmid. This PCR product and pLJC5-TMEM192-3XHA (5) were restriction digested by AgeI HF and EcoRI HF, and the 3XHA-EGFP-PEX26 sequence was subsequently cloned into pLJC5.

To create pLJC5-3XMyc-EGFP-PEX26, the 3XMyc-EGFP-PEX26 sequence was amplified by PCR using oGJR226 and oGJR225 from the pMXs-3XHA-EGFP-PEX26 plasmid. This PCR product and pLJC5-TMEM192-3XHA (5) were restriction digested by AgeI HF and EcoRI HF, and the 3XMyc-EGFP-PEX26 sequence was subsequently cloned into pLJC5.

To create the pLJC5 empty vector, oGJR237 and oGJR238 were annealed, pLJC5-TMEM192-3XHA (5) was restriction digested by AgeI HF and EcoRI HF, and the restriction-digested plasmid and annealed oligonucleotides were subsequently used to generate the pLJC5 empty vector.

For all experiments in this work, only the lentiviral pLJC5 empty vector, pLJC5-3XHA-EGFP-PEX26 construct (HA-PEROXO construct), and pLJC5-3XMyc-EGFP-PEX26 construct (Control-PEROXO construct) were used.

### Availability of Control-PEROXO and HA-PEROXO constructs

The lentiviral Control-PEROXO (Addgene #139059) and HA-PEROXO constructs (Addgene #139054) can be obtained through Addgene.

### Generation of empty vector, Control-PEROXO, and HA-PEROXO cells

Lentiviruses harboring the pLJC5 empty vector, Control-PEROXO, and HA-PEROXO constructs were generated by transfection of HEK-293T cells with the desired constructs and plasmids containing the *vsv-g* and *Δvpr* genes via standard methods. HEK-293T and T47D cells were then transduced using standard methods with viruses harboring the empty vector, Control-PEROXO, or HA-PEROXO constructs at low multiplicity of infection (10% survival after puromycin selection). Briefly, 500,000 HEK-293T or 250,000 T47D cells were plated in each well of a 6-well plate in media containing polybrene (8 μg/ml) and varying amounts of virus-containing media. Cells were centrifuged for 45 min at 774 × *g* at 37°C. 16 h later, the cells were given fresh media. The following day, cells were trypsinized and plated in media containing puromycin (1 μg/ml). After puromycin selection, cells were sorted for low EGFP levels via FACS to reduce the chances of unwanted effects from excessive amounts of the Control-PEROXO and HA-PEROXO proteins, such as mislocalization of the epitope-tagged proteins and perturbations of native cell biology. The FACS gating strategy for the Control-PEROXO and HA-PEROXO cells was to isolate a population of cells with EGFP positivity just beyond the EGFP signal distribution in the corresponding negative control.

### Cell culture

HEK-293T and T47D cells and their derivatives were maintained at 37°C and 5% CO_2_ in high glucose, GlutaMAX, sodium pyruvate, DMEM supplemented with 10% inactivated fetal calf serum, penicillin, and streptomycin. The HEK-293T and T47D cell lines were authenticated by the Duke University DNA Analysis Facility. All cells used in this study were tested for mycoplasma contamination.

### Protein extraction from whole cells

For generation of the data in Figure 2A, protein lysates were prepared in NP40 lysis buffer containing 50 mM Tris-HCl, pH 7.5, 150 mM NaCl, 1 mM EDTA, 1% NP40 and one complete mini EDTA-free protease inhibitor per 10 ml. Lysates were kept at 4°C for 15 min and then clarified by centrifugation in a microcentrifuge at 21,130 × *g* at 4°C for 10 min.

### PEROXO-IP workflow

One plate of cells (~15 million T47D cells or ~35 million HEK-293T cells) was used for an experiment such as assessing the immunoprecipitate material by immunoblotting. Four plates of HEK-293T cells (~140 million cells total) were used for each replicate for proteomics. The PEROXO-IP workflow was generally done as described previously (3) but with additional details and modifications provided here. Cells were rinsed twice with pre-chilled PBS and then scraped in 1 ml of KPBS and pelleted at 1000 × *g* for 2 min at 4°C. Cells were then resuspended in 1000 μl of KPBS, and 5 μl (equivalent to 0.5% of the total number of cells) was added to 50 μl of NP40 lysis buffer for whole-cell protein samples. NP40 lysis buffer contained 50 mM Tris-HCl, pH 7.5, 150 mM NaCl, 1 mM EDTA, 1% NP40 and one complete mini EDTA-free protease inhibitor per 10 ml. The remaining cells were gently homogenized with 25 strokes of a 2 ml homogenizer. The homogenate was then centrifuged at 1000 × *g* for 2 min at 4°C to pellet nuclei and cells while cellular organelles including peroxisomes remained in the supernatant which was incubated with 200 μl of anti-HA magnetic beads prewashed with KPBS three times. Immunoprecipitates were then incubated rocking at 4°C for 3.5 min before being gently washed three times with KPBS using a DynaMag Spin Magnet, changing tubes after the first and last wash. It takes ~10 min to have isolated peroxisomes using this workflow after cells have been homogenized. Beads with bound peroxisomes were resuspended in 50 μl ice-chilled NP40 lysis buffer to extract proteins. Protein fractions were incubated at 4°C for 10 min, and then beads were removed with the magnet. Samples were then clarified by centrifugation in a microcentrifuge at 21,130 × *g* at 4°C for 10 min. For proteomics experiments, all the steps were performed using low protein binding tubes to minimize variability between samples and maximize the recovery of proteins. Note that NP40 was used for all detergent lysis in this study because of its compatibility with our proteomic instrumentation and for the sake of consistency across all experiments. It is worth emphasizing that different cell lines can behave differently in the PEROXO-IP workflow and may require further optimization with regards to expression of the tag, amount of input material, degree of homogenization, etc.

### Immunoblotting

Protein lysates from either whole cells or immunoprecipitates were resolved by SDS-PAGE electrophoresis at 120 V. Proteins were transferred for 2 h at 45 V to PVDF membranes. Membranes were blocked with 5% nonfat dry milk in TBST (Tris-buffered saline with 0.1% Tween-20) before incubation with primary antibodies in 5% BSA in TBST at the dilutions listed in the “Antibodies” section. Membranes were washed three times in TBST before incubation with species-specific HRP-conjugated antibodies at the dilutions listed in the “Antibodies” section. Membranes were washed again three times in TBST before visualization with ECL substrate.

### Immunofluorescence

T47D cells were plated on fibronectin-coated glass coverslips in 6-well, cell-culture dishes at 300,000 cells/well. After 12 h, the coverslips were washed once in PBS and subsequently fixed using 2 ml of 4% paraformaldehyde in PBS for 15 min at room temperature. The coverslips were washed twice in 2 ml PBS and then were permeabilized using 2 ml of 0.05% Triton in PBS for 10 min at room temperature. The coverslips were washed three times in 2 ml PBS and then blocked using 2 ml Odyssey Buffer for 40 min at room temperature. The cover slips were then incubated as needed with anti-ABCD3, anti-HA, and anti-Myc primary antibodies in Odyssey Buffer for 1 h at room temperature using the dilutions indicated in the “Antibodies” section. Afterwards, the coverslips were rinsed three times in PBS and incubated as needed with 488 donkey anti-mouse and 594 donkey anti-rabbit secondary antibodies in Odyssey Buffer for 1 h at room temperature in the dark using the dilutions indicated in the “Antibodies” section. The coverslips were then washed three times with PBS and once in dH_2_O. Coverslips were mounted on slides using Vectashield containing DAPI. Images were acquired on a Zeiss AxioVert200M microscope with a 63X oil-immersion objective and a Yokogawa CSU-22 spinning-disk confocal head with a Borealis modification (Spectral Applied Research/Andor) and a Hamamatsu Orca-ER CCD camera. The MetaMorph software package (Molecular Devices) was used to control the hardware and image acquisition. Note that staining of epitope tags was done despite the presence of EGFP in the respective Control-PEROXO and HA-PEROXO proteins because cells are sorted for low EGFP levels in our methodology, thus rendering the EGFP signal weak.

### Proteomics

Sample preparation for mass spectrometry was performed as follows. Protein digestion was performed on an S-Trap Micro Spin Column (Protifi) following the manufacturer’s protocol. Briefly, samples were mixed with 5% SDS, 50 mM triethylammonium bicarbonate (TEAB) pH 7.55, 5 mM TCEP (final concentrations), heated for 10 min at 55°C, cooled to room temperature, and alkylated with 15 mM methyl methanethiosulfonate (MMTS) for 10 min at room temperature. Afterward, phosphoric acid was added to 1.2% final concentration, and an additional 6X volume of S-Trap binding buffer (90% methanol, 100 mM TEAB, pH 7.1) was added. Samples were applied to an S-Trap Micro Spin Column and centrifuged for 1 min at 4000 × *g*. The column was washed four times with S-Trap binding buffer. Sequencing-grade trypsin (0.75 μg for IP samples or 1 μg trypsin for 25 μg of whole cell sample) in 50 mM TEAB, pH 8 was added to the column and digested for 16 h at 37°C. Peptides were eluted with 40 μl each of 50 mM TEAB and then 0.2% aqueous formic acid, followed by a final elution with 35 μl 50% acetonitrile, 0.2% formic acid. The three step-wise elutions were pooled and lyophilized.

Off-line chromatographic separation was done as follows. For generation of data-dependent acquisition (DDA) spectral libraries, two separate pools consisting of either whole-cell or IP samples were made and fractionated off-line to generate diverse pools. Briefly, a first dimension of chromatographic separation with fraction collection was utilized where the labeled tryptic peptides were subjected to basic (high pH) reversed-phase high performance liquid chromatography (HPLC) with fraction collection using Shimadzu LC-20AD pumps and an FRC-10A fraction collector. The sample was loaded on a 10 cm × 2.1 mm column packed with 2.6 μm Aeris PEPTIDE XB-C18 media (Phenomenex). The gradient was isocratic 1% A buffer (20 mM ammonium formate pH 10 in water) for 1 min 150 μl min^−1^ with increasing B buffer (acetonitrile) concentrations to 16.7% B at 20.5 min, 30% B at 31 min and 45% B at 36 min. For the whole-cell samples, 2 μg of peptides from four replicates of Control-PEROXO and HA-PEROXO cells were pooled and collected as a total of 20 fractions (fx). These were combined as follows for a total of eight samples (fx1+2+19+20, fx3+10, fx4+11, fx5+12, fx6+13, fx7+14, fx8+15+17, fx9+16+18). Similarly for the IP samples, 30% of peptides from four replicates were pooled, collected as a total of 16 fractions, and combined as follows for a total of six samples (fx1+2+15+16, fx3+8, fx4+9, fx5+10, fx6+11+13, fx7+12+14). All pooled fractions were lyophilized.

Mass spectrometry was performed as follows. Samples were analyzed by HPLC using Thermo EASY-nLC 1200 pumps and autosampler and a Thermo Q Exactive HF-X Hybrid Quadrupole-Orbitrap mass spectrometer in a nano-flow configuration. Samples were loaded on a 6 cm × 100 μm column packed with 10 μm ODS-A C18 material (YMC) and washed with 12 μl total volume to trap and wash peptides. These were then eluted onto the analytical column, which was 14 cm × 75 μm PicoFrit Emitter with 15 μm tip (New Objective) self-packed with 1.7 μm Aeris C18 material (Phenomenex). For whole cell samples, the gradient was isocratic 1% A buffer (1% formic acid in water) for 1 min 300 nl min^−1^ with increasing B buffer (1% formic acid in 80% acetonitrile) concentrations to 25% B at 105 min, 40% B at 125 min, and 100% B at 126 min. For IP samples, the gradient was isocratic 6% A Buffer for 1 min 300 nl min^−1^ with increasing B buffer concentrations to 21% B at 19.5 min, 36% B at 31 min, 50% B at 36 min, and 100% B at 48.5 min. The column was washed with 100% percent B and re-equilibrated between analytical runs.

For DDA, off-line fractionated pools were analyzed and the mass spectrometer was operated in a top 20 data-dependent acquisition mode. Each DDA cycle consisted of one MS scan at 60,000 resolution, with precursor ions meeting defined criteria (AGC target: 1E6) fragmented and detected at 15,000 resolution.

For data-independent acquisition (DIA), column and gradient setup were the same as for DDA. The mass spectrometer was operated in data-independent acquisition mode, with an isolation window from 400-900 *m/z* with width of 20 *m/z* for a total of 26 bins; placement of *m/z* bins was determined using Skyline (29). Each DIA cycle consisted of two MS full scans per completion of the MS2 fragmentation of the isolation window. Ions were detected at resolution of 30,000.

DDA analysis was done as follows. Raw files of off-line fractionated pools were loaded into Proteome Discoverer 2.2 (Thermo Fisher Scientific) as fractions and analyzed using precursor-based quantification. SequestHT was used for searching peptides against the human genome database (Human_Uniprot_Sprt_Trembl_Isoforms_Release2019_01_Contams_20190205), allowing up to two missed tryptic cleavages. Tolerances were set to 10 ppm for precursor mass and 0.02 Da for fragment mass. PSM validation was performed using Percolator (30) for PSMs with deltaCn of 0.05 or less, with the target decoy database settings for high confidence at 0.01 and moderate confidence at 0.05. In Skyline (29), spectral libraries were assembled from all DDA sample search results, using only PSMs with a q-value cutoff score of 0.95 from the .msf Proteome Discoverer output file.

DIA analysis was done as follows. Raw files of the DIA-analyzed samples were loaded into ScaffoldDIA 2.0 (Proteome Software) and searched against the spectral library assembled from DDA off-line fractionated pooled IP samples and the human genome database (Human_Uniprot_Sprt_Trembl_Isoforms_Release2019_01_Contams_20190205), allowing up to two missed tryptic cleavages. Tolerances were set to 10 ppm for precursors and 0.02 Da for fragment tolerance and library fragment tolerance. Data acquisition type was set for non-overlapping windows with precursor window size of 20 Da. Only peptides with charges of +2 and +3 and length in the range 6–30 were considered. Peptides identified in each sample were filtered by Percolator (30) to achieve a maximum FDR of 0.01. Individual search results were combined and peptide identifications were assigned posterior error probabilities and filtered to an FDR threshold of 0.01 by Percolator. Peptide quantification was performed by Encyclopedia (0.9.2). For each peptide, the five highest quality fragment ions were selected for quantification. Proteins with a minimum of one identified peptide were thresholded to achieve a protein FDR threshold of 1%. See Dataset S1 for further details about the reported proteomic data.

### Statistics, analysis, and datasets

All *p*-values and corrections for multiple comparisons pertaining to proteomic analyses were calculated as described in Dataset S1. The correlation matrices and Spearman rs values in Fig. 3*B* were generated via GraphPad Prism 8. The *p*-values calculated for the correlation analyses were generated using unpaired, two-tailed *t*-tests and corrections for multiple comparisons were done using the Benjamini-Hochberg procedure via GraphPad Prism 8. The *p*-values in Fig. 3*D* were generated using unpaired, two-tailed *t*-tests via GraphPad Prism 8. The Gene Ontology (GO) Resource was used for annotating cellular compartments in this work (GO Resource accessed 2/16/2020) (31, 32). Please note that the GO Resource continues to receive updates and changes over time.

## AUTHOR CONTRIBUTIONS

W.W.C. initiated the project, developed the PEROXO-Tag, and established the PEROXO-IP workflow. G.J.R., W.W.C. designed the research with assistance from D.M.S. G.J.R. conducted most of the experimental work. W.W.C. and G.J.R. did most of the data analysis. E.S. and E.A.B. helped generate the raw proteomic data. E.A.B., E.S., W.W.C., and S.L. analyzed the raw proteomic data. G.A.W. assisted with general experimentation. W.W.C. wrote the manuscript with contributions from G.J.R. and E.A.B., and D.M.S. edited the manuscript.

## ACKNOWLEDGMENTS

We thank Caroline A. Lewis, Sze Ham Chan, Tenzin Kunchok, Izabella A. Pena, Jibril F. Kedir, all members of the Sabatini laboratory, Eric Spooner, and Hoi See Tsao for helpful suggestions. We thank Destiny Tolliver and all other members of the Boston Combined Residency Program for their assistance. Certain elements of figures were created with BioRender.com. G.J.R. is supported by a fellowship from the US NIH (F31GM129905-01). E.A.B. is a Robert Black Fellow of the Damon Runyon Cancer Research Foundation (DRG-2365-19). G.A.W. is presently a Howard Hughes Medical Institute Jane Coffin Childs Fellow. This work was supported by grants from the US NIH (R01AI144369 to S.L.; R01CA103866, R01CA129105, and R37AI047389 to D.M.S) and the Department of Defense (W81XWH-15-1-0230 to D.M.S.). W.W.C. is supported by the Boston Combined Residency Program and Boston Children’s Hospital. D.M.S. is an investigator of the Howard Hughes Medical Institute and an American Cancer Society Research Professor.

